# Targeting CXCR1 and CXCR2 to overcome radiotherapy resistance in PTEN-deficient prostate carcinoma

**DOI:** 10.1101/2020.03.16.993394

**Authors:** Chris W.D. Armstrong, Jonathan A. Coulter, Chee Wee Ong, Pamela J. Maxwell, Steven Walker, Karl T. Butterworth, Oksana Lyubomska, Silvia Berlingeri, Rebecca Gallagher, Joe M. O’Sullivan, Suneil Jain, Ian G. Mills, Kevin M Prise, Robert G. Bristow, Melissa J. LaBonte, David J.J. Waugh

**Author notes:** These authors contributed equally to this manuscript (Melissa J. LaBonte, David J.J. Waugh). Correspondence: Dr Melissa LaBonte, Centre for Cancer Research and Cell Biology, Queen’s University Belfast, 97 Lisburn Road, Belfast BT9 7AE, Northern Ireland.

## Abstract

Functional impairment of the tumour-suppressor *PTEN* is common in primary-prostate cancer and has been linked to relapse post-radiotherapy (RT). Pre-clinical modelling supports elevated CXC-chemokine signaling as a critical mediator of *PTEN*-depleted disease progression and therapeutic resistance. We assessed the correlation of *PTEN*-deficiency with CXC-chemokine signaling and its association with clinical outcomes. Gene expression analysis characterized a *PTEN*^LOW^/CXCR1^HIGH^/CXCR2^HIGH^ cluster of tumors that associates with earlier time-to-biochemical recurrence (HR 5.87 and HR 2.65 respectively) and development of systemic metastasis (HR 3.51). *In vitro*, CXCL-signaling was further amplified following exposure of *PTEN*-deficient prostate cancer cell lines to ionizing radiation (IR). Inhibition of CXCR1/2-signaling in *PTEN-*depleted cell-based models increased IR-sensitivity. *In vivo*, administration of a CXCR1/2-targeted pepducin (x1/2pal-i3), or CXCR2-specific antagonist (AZD5069), in combination with IR to *PTEN*-deficient xenografts attenuated tumor growth and progression compared to control or IR alone. Post-mortem analysis confirmed that x1/2pal-i3 administration attenuated IR-induced CXCL-signaling and anti-apoptotic protein expression. Interventions targeting CXC-chemokine signaling may provide an effective strategy to combine with radiotherapy, in both locally-advanced and oligometastatic-prostate cancers, with known presence of *PTEN*-deficient foci.

## INTRODUCTION

External beam radiotherapy constitutes a principal treatment modality for organ-confined prostate cancer (CaP) (Resnick, Koyama et al., 2013). Although the majority of tumors respond favourably, over a third of patients will experience relapse post-radiotherapy (RT), which has been attributed to intrinsic radioresistance of tumor cells, release and signaling of stroma-derived survival factors, or presence of occult micro-metastases at the time of diagnoses (Barker, Paget et al., 2015, Darwish & Raj, 2012, Miyake, Tanaka et al., 2014). The cellular response of tumor cells to DNA-damage treatment can be related to an underlying genetic background and can itself markedly alter gene expression to effect differential phenotypic behaviour (Kumareswaran, Ludkovski et al., 2012, Taiakina, Dal Pra et al., 2014). Crucially, RT is no longer restricted to use in the treatment of local disease but is becoming a viable therapeutic option for advanced disease, with clinical trials currently evaluating the potential use of stereotactic RT to treat oligometastatic CaP (Muldermans, Romak et al., 2016, Ost, Jereczek-Fossa et al., 2016, Patel, Chaw et al., 2019). The extended deployment of radiation as an intervention across the continuum of the clinical landscape accentuates the requirement to optimize this treatment modality.

Improving the effectiveness of RT has typically followed two distinct pathways: [1] altering standard treatment protocols with the intent to boost the overall radiation dosage delivered to the tumor, including the recent introduction of hypo-fractionated dose scheduling (Arcangeli, Saracino et al., 2017, Catton, Lukka et al., 2017, Dearnaley, Syndikus et al., 2016); and [2] the use of genetic markers to stratify patients and/or progress combination-targeted therapy to attenuate associated survival mechanisms adopted by cells as a mechanism of resistance to radiation-induced cell death. Zafarana and colleagues originally reported allelic loss of the tumour suppressor *PTEN* and allelic gain of *c-MYC* as prognostic factors for relapse following RT (Zafarana, Ishkanian et al., 2012b). Identifying the critical signaling pathways that underpin and confer *PTEN*-mediated resistance is essential to defining actionable combinatorial drug-radiotherapy treatment approaches that may be employed to improve radiotherapy response in future patients (Armstrong, Maxwell et al., 2016, McCabe, Hanna et al., 2015).

*PTEN*, a potent negative regulator of the PI3K-Akt signaling axis, is deleted or mutated in approximately 30% of men with localized CaP and in over 60% of patients exhibiting metastatic progression (Phin, Moore et al., 2013, Suzuki, Freije et al., 1998). Moreover, impairment of PTEN function is associated with clinico-pathological features of aggressive and treatment-resistant prostate carcinoma (Ferraldeschi, Nava Rodrigues et al., 2015, Khemlina, Ikeda et al., 2015, Zafarana, Ishkanian et al., 2012a). Our initial studies confirmed the increased expression of CXCL8 and its two receptors CXCR1 and CXCR2 in the tumor epithelium of human CaP (Murphy, McGurk et al., 2005). Our subsequent studies in human CaP cell lines and genetically-engineered mouse models associated the elevated expression of this chemokine signaling pathway with *PTEN*-loss (Maxwell, Coulter et al., 2013). Intrinsic CXCL8-signaling underpins prostate cancer cell survival through the activation of AR, HIF-1 and NF-κB transcription factors, and increases expression of anti-apoptotic proteins, including members of the Bcl-family (Maxwell, Gallagher et al., 2007). In addition to underpinning resistance to AR-targeted therapy, induction of CXCL8 signaling modulates the sensitivity of prostate cancer cells to several novel molecular targeted therapies and chemotherapeutic agents, including to oxaliplatin which induces DNA double-strand breaks in CaP cells (Wilson, Purcell et al., 2008). Thus, we adopted the hypothesis that exposure to ionizing radiation (IR) would similarly induce chemokine-signaling and that this would have a profound impact in modulating the sensitivity of *PTEN*-deficient tumors to radiation.

The objective of this comprehensive study, employing clinical datasets and established *in vitro* and *in vivo* models, was to characterize whether stress-induced potentiation of CXCR1/2-signaling may underpin the adverse response of *PTEN*-deficient prostate cancer to radiation and to potentially explain biological mechanisms related to increased clinical relapse reported in *PTEN*-deficient tumors.

## MATERIALS AND METHODS

### Cell culture

Authenticated DU145, LNCaP, C42, C42B, PC3 and 22Rv1 CaP cells were obtained from ATCC. DU145 cells were manipulated as previously described to generate isogenic PTEN-expressing NT01 cells and PTEN-deficient sh11.02 cells (Maxwell et al., 2013). PC3 were manipulated as previously described so that PTEN-expression can be reconstituted following exposure to tetracycline (Maxwell et al., 2013). PTEN-depleted 22RV1 cells were generated following lentiviral transfection of HuSh-29 pre-designed PTEN shRNA pGFP-V-RS constructs (Origene, Rockville, MD, USA), and selected under puromycin selection pressure at a final concentration of 0.5 μg/ml. Cell line authenticity was confirmed by STR genotyping (July 2019) and mycoplasma testing was performed every 4-6 weeks (MycoAlert, Lonza).

### ELISA

Cells were plated into six-well plates at a density of 5 × 10^5^ cells per well and allowed to adhere overnight. After 24H, cells were irradiated and media samples collected at various timepoints. Cell counts were performed at each timepoint. CXCL8 ELISA experiments were performed using DuoSet® ELISA Development Kits (R&D Systems) according to manufacturer’s instructions. CXCL8 secretion was normalized to cell count to correct for differences in confluency.

### Immunoblotting

Whole cell lysates were prepared, resolved and blotted as previously described (Maxwell et al., 2013). Membranes were probed with primary antibodies at 4°C overnight. Primary antibody information can be found in Supplementary Table 1. Following three TBST washes, membranes were incubated with the appropriate horseradish peroxidase (HRP)-labeled secondary antibody (1:5000 dilution; GE Healthcare UK Ltd, UK). Protein bands were detected using enhanced chemiluminescence (Luminata Crescendo, Merck Millipore). Membranes were re-probed with β-Actin antibody to ensure equal loading.

### Quantitative real-time PCR (qRT-PCR)

Total RNA was collected and isolated as previously described (Maxwell et al., 2013). Quantitative real-time polymerase chain reaction (qRT-PCR) was performed using pre-validated RealTime ready custom assays for *CXCR1, CXCR2, CXCL8, and BCL2* used in combination with FastStart TaqMan® Probe Master solutions (Roche Diagnostics, Sussex, UK). Individual sample mRNA levels were analysed in triplicate in a 96-well plate using an LC480 light cycler instrument (Roche Diagnostics). Gene expression levels were normalised against 18S.

### siRNA

siRNA transfections for CXCR1 and CXCR2 oligonucleotides (Dharmacon, Lafayette, CO, USA) were carried out using Lipofectamine® RNAiMAX Transfection Reagent (Life Technologies, Paisley, UK) when cells reached 60-70% confluence. Briefly, for a p90 petri dish, 10 μl RNAiMAX was combined with 25 nM pooled CXCR1 and CXCR2 oligonucleotides (12.5 nM CXCR1/ 12.5 nM CXCR2) and added in a drop-wise fashion to 2 mL Opti-MEM. Transfection complexes were then incubated at room temperature for 20 min, after which the siRNA complexes were added to 8 mL of complete medium. Cells were then maintained at 37°C for 48H. Non-targeting sequences were used at the same concentration (25 nM) as the total combined CXCR1 and CXCR2 siRNA sequences.

### Flow cytometry

Cells were seeded at a density of 5×10^4^ per well in six-well plate and left to adhere overnight. Cells were then transfected with the appropriate control or siRNA oligonucleotides and returned to the incubator. After 72H, all groups received either a 3 Gy radiation dose or sham irradiation. Cells were analyzed 72H post-radiation treatment. Whole culture medium was collected and pooled with the trypsinized cells, then centrifuged at 1500 rpm. Cell pellets were resuspended in 100 μL of 1X binding buffer. Annexin V (Life Technologies, Paisley, UK) antibody (5 μL) was added to each sample along with 5μL of propidium iodide (PI) stain (50 μg/mL). Samples were then incubated in the dark, at room temperature for 15 min. After incubation, 320 μL of 1X binding buffer was added to each sample before analysis on the EPICS XL flow cytometer (Beckman Coulter, Buckinghamshire, UK).

### Clonogenic assays

Reverse clonogenic assays were performed as follows. In CXCR1/2-targeting experiments, cells were transfected, irradiated 48 h post-transfection and reseeded to assess colony-forming ability. In PC3-PTEN cells, transfections were performed 24 h prior to *PTEN*-reconstitution using tetracycline (1 μg/ml). Surviving fractions (SF) were calculated relative to non-irradiated cells and fitted using a linear quadratic function (S = exp(-αD -βD2)) using least-squares regression (Prism 7.0; GraphPad Software, CA). Area under-the-curve (AUC) representing the mean inactivation dose (MID) was obtained and dose enhancement factor (DEF) calculated by dividing the MID of the CXCR1/2-depleted cells by that of non-targeting siRNA treated cells.

### PC3 and DU145 xenograft tumor growth delay models

CXCR1/2 blocking pepducins (CXCR1/2 x1/2pal-i3 pepducin - sequence pal-RTLFKAHMGQKHR-NH2; non-targeting x1/2pal-con – sequence pal-TRFLAKMHQGHKR-NH2) which target the highly conserved third intracellular loop were used to block chemokine receptor-mediated signaling *in vivo*. Pepducin targeting and efficacy was confirmed *in vitro* prior to *in vivo* use. PC3, NT01 or sh11.02 cells (2 × 10^6^ in PBS) were implanted by intradermal injection on the rear dorsum of BALB/c SCID mice (Envigo). Animals with palpable tumors (100mm^3^) were randomized to treatment groups (N=7/group): no treatment; x1/2pal-con, x1/2pal-i3, 3 Gy, x1/2pal-con + 3 Gy and x1/2pal-i3 + 3 Gy. Pepducins, reconstituted in PBS, were administered by intra-tumoral injection (2 mg/kg) on days 1, 2, 3, 4 and 5. Radiation was administered on day 3. Animals were restrained in a perspex jig and protected from non-target radiation damage using lead shielding. Radiation treatments were delivered as two parallel-opposed fields using a Precision X-RAD 225. Due to the increased radiation sensitivity of DU145 cells, radiation dose was reduced to 2 Gy. Tumor dimensions were measured using calipers and tumor volumes calculated using the formula (width^2^ x length)/2. Tumor and weight measurements were performed every Monday, Wednesday and Friday for the duration of the study. Animals were culled when the tumor volume quadrupled (400mm^3^). Additional mice (N=4/group) were culled on study day 5 to enable collection of tumors for pharmacokinetic analysis. Tumors were cut into two halves and stored in either formalin or liquid nitrogen.

### C4-2 xenograft model

SCID male mice (7 weeks old) were obtained from Envigo. C4-2 cells were injected subcutaneously, 1 × 10^7^ cells in 100 uL PBS:Matrigel (50:50) in the right flank. Once palpable tumors formed, tumour volume was measured by digital calipers, as described above. When tumors reached 100mm^3^, mice were randomized to treatment groups (N=8/group): vehicle control, AZD5069, 3 Gy, or AZD5069+3 Gy. AZD5069 was prepared in 1X PBS/0.1% Tween-20 and administered by oral gavage at 2 mg/kg daily. Radiation was administered on day 3, as described above. Tumor and weight measurements were performed 3x weekly for the duration of the study. Animals were culled when the tumor volume reached maximum (1000mm^3^). Additional mice (N=4/group) were culled on study day 5 to enable collection of tumors for pharmacokinetic analysis. Tumors were cut into two halves and stored in either formalin or liquid nitrogen.

### 53BP1 foci immunofluorescence

Cells (1 × 10^4^) were seeded onto 4-well chamber slides (Fisher Scientific, Leicestershire, UK) and left overnight to adhere. For *PTEN*-depleted damage repair studies, cells were irradiated with 1 Gy (to prevent damage saturation) and analysed at indicated time points ranging for 0.5 to 24H post-radiation to observe repair kinetics. For CXCR1/CXCR2 knockdown experiments, transfections were performed 72H prior to irradiation. Cells were fixed 4H post-radiation treatment with 50% methanol: 50% acetone, and permeabilised in 0.5% triton X-100 (Sigma, Dorset, UK). After an overnight incubation in blocking buffer (PBS, 0.1% triton X-100, 5% FCS, 0.2% milk), cells were incubated with rabbit anti-53BP1 antibody (Abcam, Cambridge, UK) at 1:2000 concentration and incubated with secondary anti-rabbit Alexa fluor-488 (Life Technologies, Paisley, UK). Nuclei were counterstained with 4,6-diamidino-2-phenylin-dole (0.1 mg/mL). Foci were counted using a fluorescence microscope (Zeiss Axiovert 200M, UK); typically, 100 cells were counted per treatment condition.

### Immunohistochemistry

Sections were cut from tumor tissue blocks for H&E and IHC. The initial section was used for H&E staining to assess histology and appropriate tumor content for subsequent IHC localisation and analysis. Sections for IHC were cut at 4 µM on a rotary microtome, dried at 37°C overnight and used for IHC tests, performed on an automated immunostainer (Leica BOND-MAX™). Validated and optimised protocols, used in local diagnostics, were selected for each biomarker with inclusion of carefully selected control tissues during antibody application. Antigen binding sites were detected with a polymer-based detection system (Bond cat no. DS 9800). All sections were visualized with 3,3’-diaminobenzidine (DAB), counterstained in haematoxylin and mounted in DPX. Biomarker conditions were as follows: Ki67 (clone MM1 DO-7 mouse monoclonal antibody, Leica) was used at a 1:200 dilution with epitope retrieval solution 2 pre-treatment for 30 min. Bcl-2 (clone 3.1 mouse monoclonal antibody, Leica) was used at a 1:100 dilution with epitope retrieval solution 2 pre-treatment for 20 min.

### Statistical Analysis

Unless otherwise stated, experiments were performed in triplicate and results expressed as mean±standard error (SE). Data was analyzed using two-tailed unpaired t-test or two-way ANOVA (clonogenic survival curves) with p-value of <0.05 considered to be statistically significant.

## RESULTS

### Correlation of PTEN and CXC-chemokine signaling to clinical parameters of progression

*PTEN*-deficiency increases *CXCL8*, *CXCR1* and *CXCR2* gene expression in human CaP cell lines *in vitro* and that of the orthologous pathway in the prostate epithelium of *PTEN*^+/-^ mice (Maxwell et al., 2013). We conducted *in silico* analysis of publicly-available CaP datasets to confirm this association in human prostate cancer and its association with clinical outcomes. Our initial analysis was conducted using the MSKCC radical prostatectomy cohort, focusing on the 140 patients with relevant clinical follow-up data (Taylor, Schultz et al., 2010). This dataset derived from radically-resected tumors demonstrated that *PTEN^LOW^* (*p*=0.0014), CXCR1^HIGH^ (*p*=0.017) and CXCR2^HIGH^ (*p*=0.035) expression each independently correlated with accelerated biochemical recurrence (BCR) (Figure 1A). We observed that the *PTEN*^LOW^, CXCR1/2^HIGH^ tumors formed a distinct cluster or sub-group constituting 74 (52.9%) of the 140 tumors represented in this resection cohort (Figure 1B). Kaplan-Meier analysis revealed the PTEN^LOW^/ CXCR1/2^HIGH^ cluster was associated with a highly significant reduction in time to BCR (*P*<0.001; HR: 5.87) (Figure 1C).

**Figure 1.**
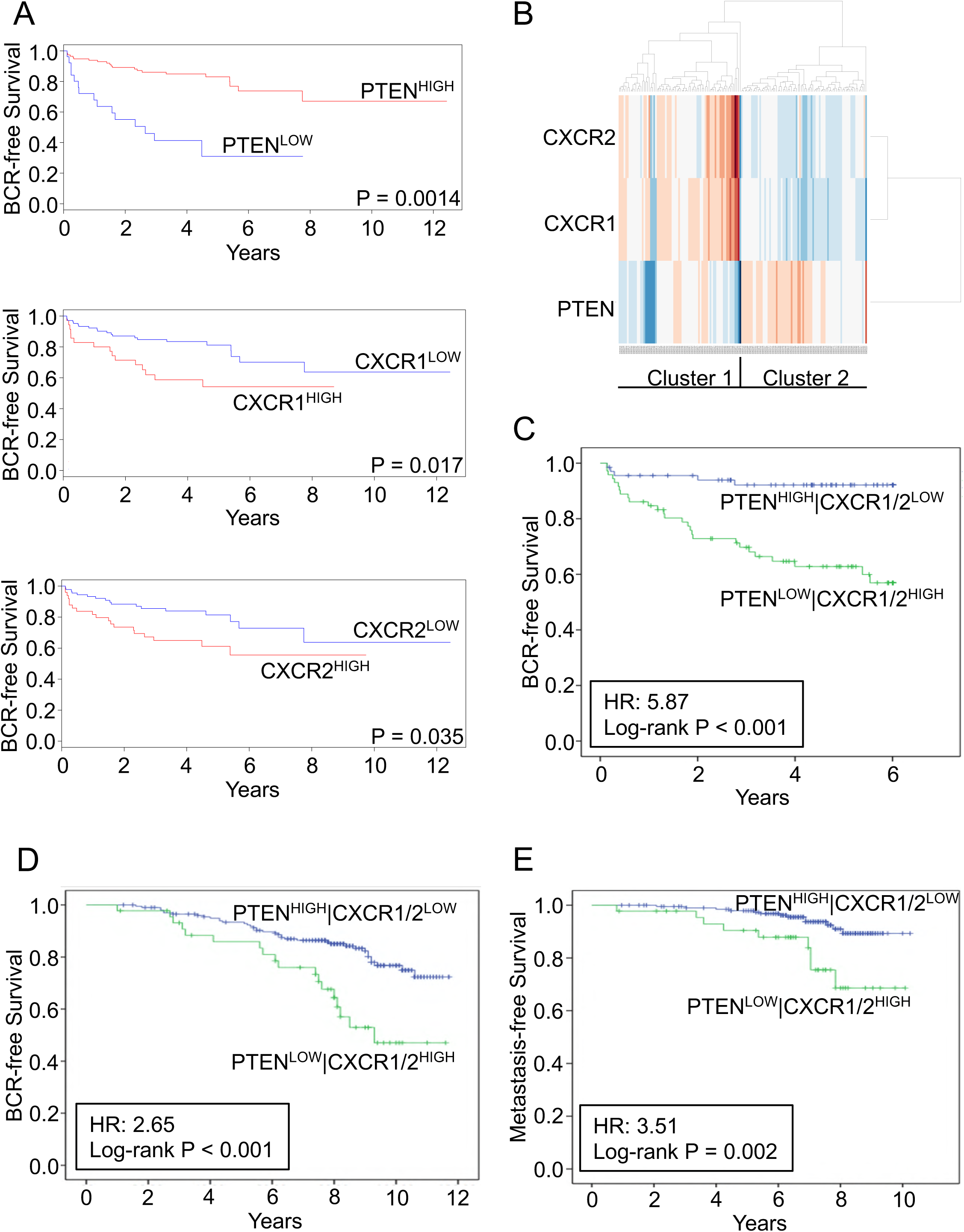
*PTEN^LOW^*, *CXCR1^HIGH^* and *CXCR2^HIGH^* tumors are associated with poor prognosis. **A** Kaplan-Meier curves examining the relevance of *PTEN*, *CXCR1* and *CXCR2* expression independently in the Taylor (MSKCC) dataset for biochemical recurrence (BCR). High and low expression was of each gene was determined based on differences from the median threshold (*PTEN*: 8.63; *CXCR1*: 6.27; *CXCR2*: 6.59). Sample sizes were as follows: *PTEN^HIGH^* (N=114); *PTEN^LOW^* (N=26); *CXCR1^HIGH^* (N=35); *CXCR1^LOW^* (N=105); *CXCR2^HIGH^* (N=50); *CXCR2^LOW^* (N=90). **B** Cluster analysis of *PTEN*, *CXCR1* and *CXCR2* in the MSKCC dataset. **C** Kaplan-Meier survival curves in 140 patients from the MSKCC dataset in relation to sample clustering by three genes (*PTEN*, *CXCR1* and *CXCR2*). Sample sizes were as follows: *PTEN^HIGH^|CXCR1/2^LOW^* (n=66); *PTEN^LOW^|CXCR1/2^HIGH^* (N=74). **D-E** Analysis of the FASTMAN retrospective radiotherapy patient sample cohort (N=248) showing the impact of previously defined clusters on time to recurrence and time to metastasis respectively. Sample sizes were as follows: *PTEN^HIGH^|CXCR1/2^LOW^* (N=203); *PTEN^LOW^|CXCR1/2^HIGH^* (N=45). Data information: Significant differences were determined by log-rank test. Abbreviations: BCR, biochemical recurrence; HR, hazard ratio.

Further analysis was performed on a second dataset to determine the direct relevance of this gene cluster with respect to radiotherapy response. Analysis was conducted on a transcriptomic profile derived from the FASTMAN retrospective radiotherapy patient sample cohort (Jain, Lyons et al., 2018). This cohort of 248 diagnostic biopsy samples has a median follow-up data of >100 months. Kaplan-Meier analysis confirmed that *PTEN*^LOW^, CXCR1/2^HIGH^ tumors were associated with a significantly reduced time to BCR (Figure 1D; *p*<0.001; HR: 2.65) and importantly, with the development of metastasis (Figure 1E; *p*=0.002; HR: 3.51) after RT treatment. Taken together these results establish the clinical relevance of the PTEN^LOW^/ CXCR1/2^HIGH^ cluster in two distinct cohorts, and furthermore, reveal the downstream significance of this biology in locally-advanced prostate cancer to adverse outcomes after both surgery or radiotherapy interventions.

### Expression of PTEN modulates radiation-induced CXCL-chemokine signalling

Radiotherapy is a major treatment modality for locally-advanced and increasingly for oligometastatic prostate cancer. Experiments were therefore performed across a range of CaP cell models representative of different androgen sensitivity, different metastatic potential and exhibiting differential expression of PTEN: we used *PTEN*-null LNCaP and LNCaP-derived C4-2 and C4-2B cells and isogenic PTEN-expressing and -null cell lines (DU145, PC3 and 22Rv1) (Maxwell et al., 2013). Their PTEN-expression and resulting downstream PI3K-AKT signaling axis activity were confirmed by immunoblot analysis for PTEN and phosphorylation status of AKT^S473^ (Figure 2A).

**Figure 2.**
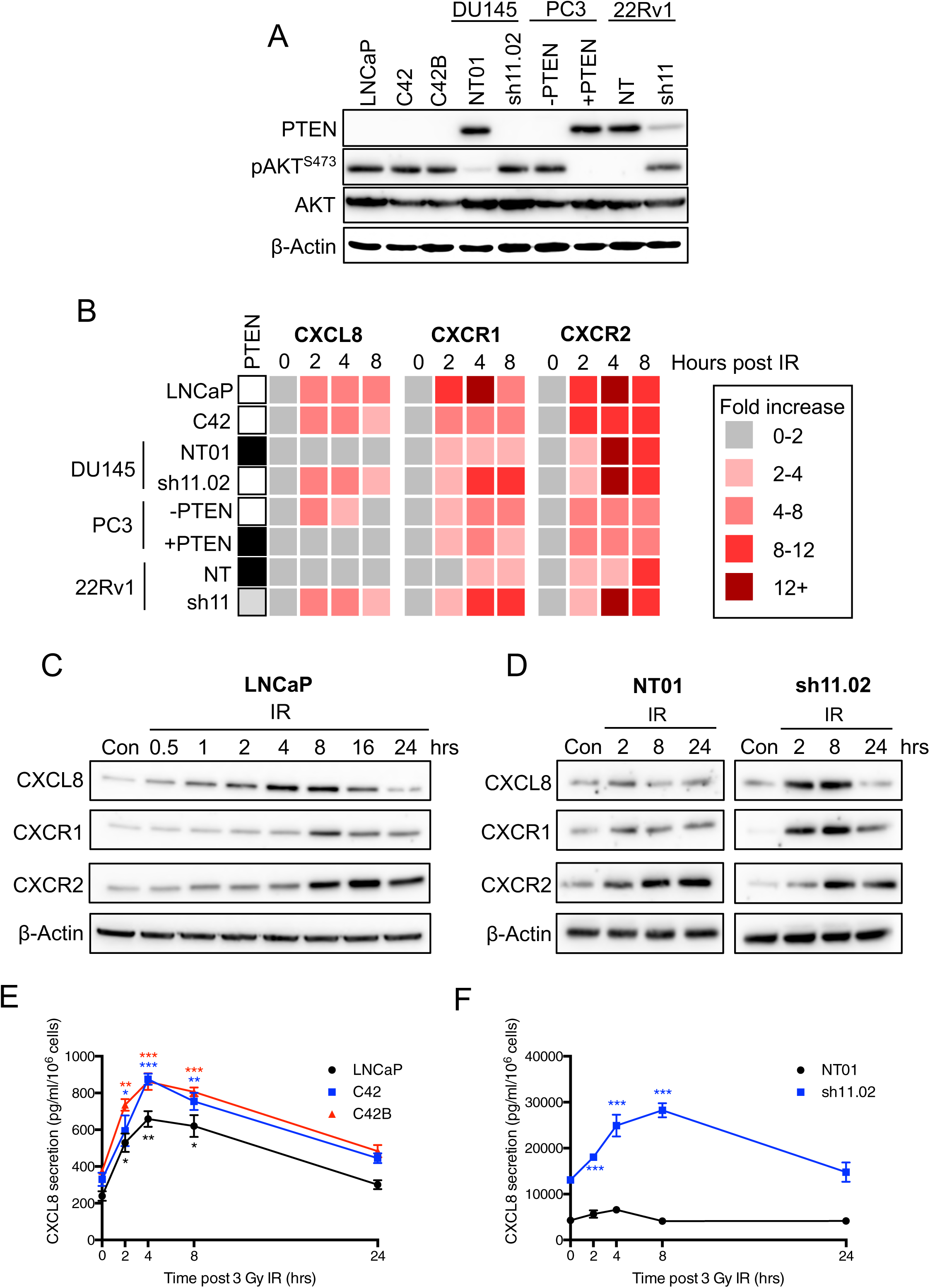
Ionizing radiation induces CXC-chemokine expression and secretion in *PTEN*-deficient prostate cancer models. **A** Immunoblots showing expression of PTEN, pAKT (Ser473) and AKT in a panel of prostate cancer cell models. PTEN expression was depleted in DU145 and 22Rv1 cells using a lentiviral-based protocol. PTEN expression was reconstituted in PC3 cells under the control of a tetracycline-inducible promoter. Equal loading was confirmed by re-probing blots for β-Actin. **B** Heatmap showing fold-increase of *CXCL8*, *CXCR1* and *CXCR2* gene expression from 0-8H following exposure to 3 Gy IR. PTEN expression of each cell line is indicated on the left-panel (white: PTEN-null; black: PTEN-expressing; grey: partial PTEN depletion). **C-D** Immunoblots showing expression of CXCL8, CXCR1 and CXCR2 at 0-24 hrs following treatment LNCaP, NT01 and sh11.02 cells with 3 Gy IR. Equal loading was confirmed by re-probing blots for β-Actin. **E** Graph showing CXCL8 secretion from LNCaP, C4-2 and C4-2B cells from 0-24 hrs following treatment with 3 Gy IR. **F** Graph showing CXCL8 secretion from NT01 and sh11.02 cells from 0-24 hrs following treatment with 3 Gy IR. Data information: All data presented is a representation of N=3 independent experiments. (E-F) Statistical analysis is a comparison of each time point to the 0 hr control within the same cell line (*p*<0.05*; *p*<0.01**; *p*<0.001***).

The role of radiation in modulating CXC-chemokine signaling was investigated using PTEN-expressing and -null CaP cells, exposed to clinically-relevant doses of radiation (2-3 Gy). qRT-PCR analysis was used to quantify alterations in *CXCL8*, *CXCR1* and *CXCR2* gene expression (Figure 2B). While *CXCL8* expression remained unchanged in cell lines that retained sufficient levels of *PTEN* (DU145-NT01, PC3-PTEN and 22Rv1-NT), PTEN-deficient cells demonstrated an increase in *CXCL8* mRNA 2H post-radiation that was sustained out to 8H. However, treatment with IR induced the expression of chemokine receptors *CXCR1* and *CXCR2* in tumor cells independent of *PTEN* status (Figure 2B).

Similar experiments were performed to assess the impact of IR on CXC-chemokine protein expression using LNCaP and DU145-NT01 and sh11.02 cells. Treatment with 3 Gy increased CXCL8 expression in *PTEN*-null LNCaP and *PTEN*-depleted sh11.02 cells; but did not modulate expression in *PTEN*-expressing NT01 cells. Confirming our qRT-PCR analysis, IR induced expression of CXCR1 and CXCR2 at the protein level was independent of intrinsic PTEN status (Figures 2C and 2D).

We further assessed the effect of IR on CXCL8 secretion. Exposure to IR induced a 3-4-fold increase in the secretion of this chemokine from *PTEN*-null LNCaP, C4-2 and C4-2B cells. This was evident 2H post-IR exposure, with levels returning to baseline within 24H (Figure 2E). Similarly, in PTEN-expressing DU145-NT01 cells and PTEN-depleted DU145-sh11.02 cells, treatment with 3 Gy resulted in significantly increased CXCL8 secretion in sh11.02 cells, but had no effect in NT01 cells (Figure 2F). Interestingly, baseline secretion was already 3.06-fold greater (*p*<0.0001) in the PTEN-depleted sh11.02 cells (Figure 2F). Crucially, we observed these increases in CXCL8 secretion across androgen-dependent and -independent models signifying the importance of this biology across the disease spectrum.

### Inhibition of CXCR1/2-signaling promotes PTEN-dependent radio-sensitization

CXCR1- and CXCR2-targeted siRNA was used to knockdown expression of both receptors in order to assess the survival advantage afforded by hyperactive CXC-chemokine signaling. Validation of these siRNAs confirmed their ability to successfully reduce expression of the respective receptors by more than 90% in LNCaP and C4-2 cells (Figure 3A). Knockdown of both receptors was also validated in PTEN-expressing DU145 NT01 cells and PTEN-depleted DU145 sh11.02 cells (Figure 3B), providing a range of experimental models to evaluate the impact of targeting CXCL8/CXCR-signaling.

**Figure 3.**
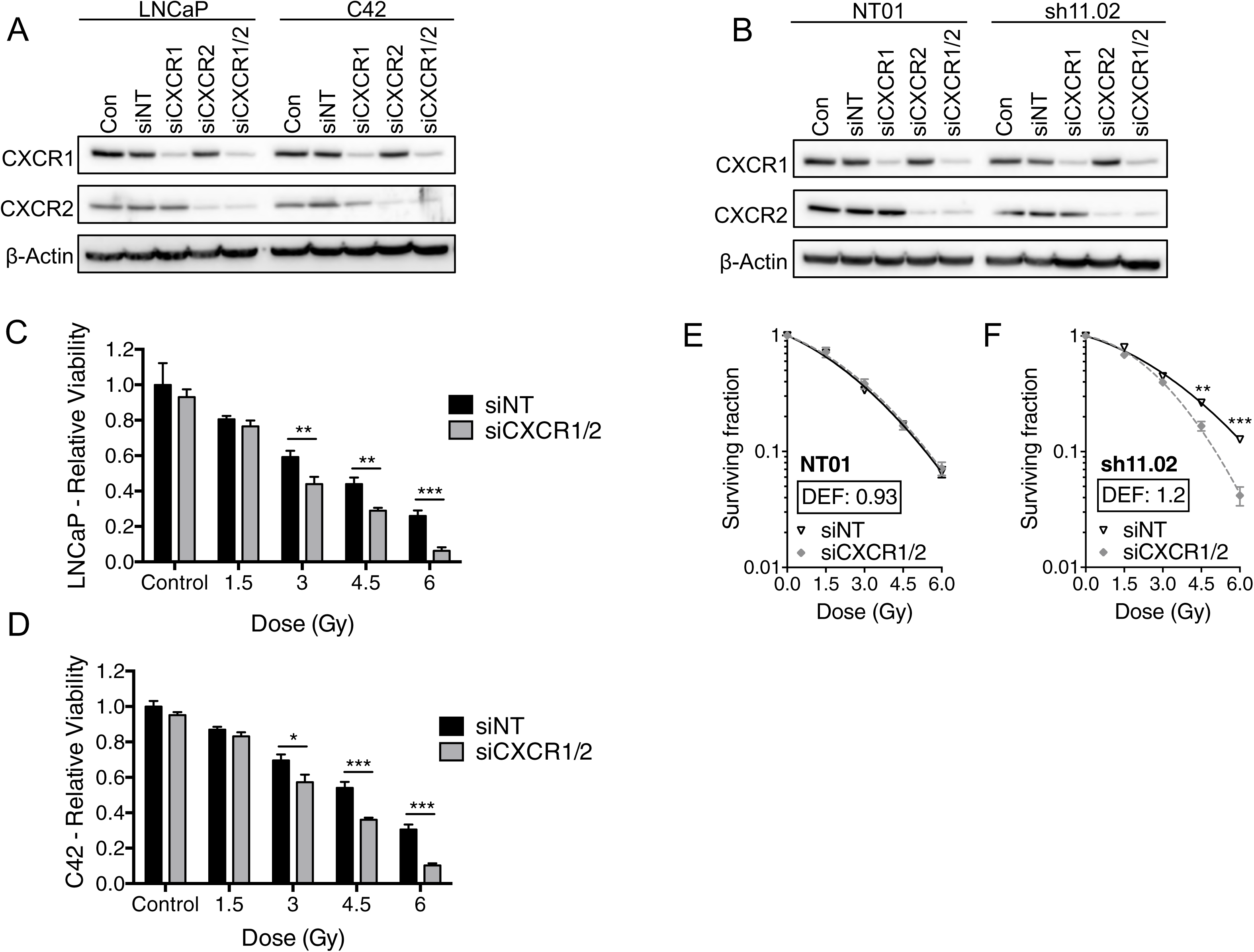
Radiosensitivity of *PTEN*-deficient prostate cancer via CXCR1/2 knockdown. **A-B** Immunoblots showing CXCR1 and CXCR2 expression following treatment with non-targeting siRNA (siNT), CXCR1 siRNA (siCXCR1), CXCR2 siRNA (siCXCR2) or a combination of both CXCR1 and CXCR2 siRNA (siCXCR1/2). Final siRNA concentration was 25 nM. Equal loading was confirmed by re-probing blots for β-Actin. **C-D** Bar charts showing relative viability of LNCaP and C42 cells respectively as determine by Alamar Blue assay. Cells were pre-treated with 25nM non-targeting siRNA (siNT) or CXCR1 and CXCR2 siRNA (siCXCR1/2) for 48H prior to treatment with IR (0-6 Gy dose range). Samples were analyzed 7 days following IR treatment. **E-F** Clonogenic survival curves generated from NT01 and sh11.02 cells respectively. Cells were pre-treated with 25nM non-targeting siRNA (siNT) or CXCR1 and CXCR2 siRNA (siCXCR1/2) for 48H prior to IR treatment. Data information: All data presented is a representation of N=3 independent experiments. All survival curves are fitted to a linear quadratic model and Dose Enhancement Factor (DEF) was calculated using the mean inactivation dose based on the area under the curve at a surviving fraction of 0.1. Statistically significant differences were determined using t-tests (*p*<0.05*; *p*<0.01**; *p*<0.001***).

The impact of *PTEN* depletion and/or repression of CXCL8-signaling on cell proliferation was determined initially by cell growth curve analysis. Knockdown of CXCR1 and CXCR2 alone had limited impact on the cell proliferation kinetics of either the PTEN-expressing DU145 NT01 or PTEN-depleted sh11.02 cells. However, siRNA-mediated suppression of CXCR1/2 enhanced the effects of IR in both the NT01 and sh11.02 cells, increasing the observed cell doubling time by 2.2-fold and >3-fold, respectively (Supplementary Table 2).

Due to the inherently poor colony forming ability of LNCaP cells, we used an Alamar Blue assay to assess the impact of CXCR1/2 knockdown upon the viability of these cells at 7 days post-radiation. Compared to cells treated with non-targeting siRNA, knockdown of both receptors significantly decreased cell viability in LNCaP and C4-2 cells once exposed to doses >3 Gy (Figures 3C-D).

Cells derived from the DU145 lineage have excellent colony forming ability and so clonogenic survival assays were used to confirm the survival advantage associated with sustained CXCR1/2-signaling. Knockdown of these receptors in PTEN-expressing NT01 cells resulted in no significant difference in survival following exposure to IR (DEF=0.93; Figure 3E). However, siRNA-mediated CXCR1/2 knockdown in *PTEN*-depleted sh11.02 cells increased sensitivity to IR with a calculated dose enhancement factor (DEF) of 1.2 (Figure 3F). Conversely, we employed a PTEN-inducible PC3 cell model, whereby the radiosensitizing effect of CXCR1/2-siRNA detected in the PTEN-null parental cell line was completely ablated following reconstitution of PTEN expression (Supplementary Figure 1).

Combined, these results confirm that a CXCR1/2-targeted therapeutic approach can drive radiosensitivity in *PTEN*-depleted prostate cancer models.

### PTEN depletion combined with knockdown of CXCR1/2 impairs tumor cell proliferation and promotes apoptosis

We have previously shown that CXCR1/2-mediated signaling results in upregulation of the anti-apoptotic, pro-survival protein, Bcl-2, suggesting that CXCR1/2 knockdown may confer radiosensitivity through modulation of the apoptotic pathway (Maxwell et al., 2013). Immunoblotting confirmed that exposure to IR induced Bcl-2 expression in androgen-dependent LNCaP cells. We also observed increased JAK2^Y1007/1008^ and STAT3^Y705^ phosphorylation providing a potential mechanism for potentiation of anti-apoptotic pathways (Dai, Wang et al.). However, knockdown of CXCR1/2 prior to IR treatment repressed the ability of LNCaP cells to induce JAK/STAT phosphorylation and subsequent Bcl-2 expression (Figure 4A). This coincided with increased detection of cleaved-PARP and -caspase 9, confirming the induction of apoptosis. Similar immunoblotting profiles were observed when experiments were repeated in the androgen-independent *PTEN*-deficient DU145 sh11.02 cells (Figure 4B).

**Figure 4.**
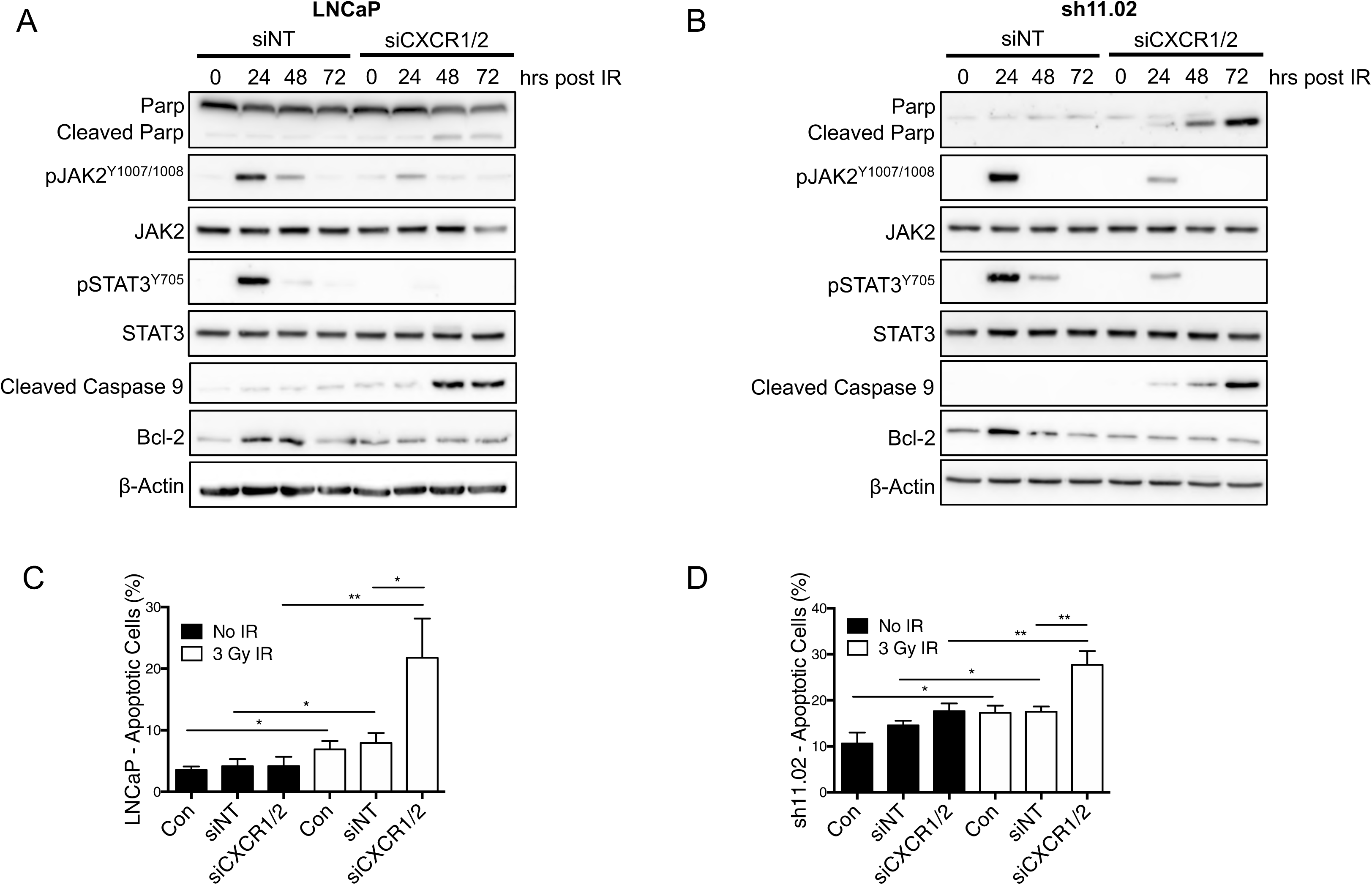
Knockdown of CXCR1 and CXCR2 enables the induction of apoptosis following treatment with radiation. **A-B** Immunoblots examining expression of an apoptotic protein panel in LNCaP and DU145-sh11.02 cells respectively. Cells were pre-treated with 25nM non-targeting siRNA (siNT) or CXCR1 and CXCR2 siRNA (siCXCR1/2) for 48H prior to treatment with 3 Gy IR. Equal loading was confirmed by re-probing blots for β-Actin. **C-D** Bar charts summarizing Annexin V/PI flow cytometry analysis of LNCaP and sh11.02 cells respectively. Cells were pre-treated with 25nM non-targeting siRNA (siNT) or CXCR1 and CXCR2 siRNA (siCXCR1/2) for 48H prior to treatment with 3 Gy IR. Cells were analyzed 72H following IR treatment alongside non-irradiated control cells (black bars). Data information: All data presented is a representation of N=3 independent experiments. Statistically significant differences were determined using t-tests (*p*<0.05*; *p*<0.01**; *p*<0.001***).

Increased apoptotic signaling was validated by analysis of apoptotic fractions using Annexin V/PI-staining protocols in LNCaP and DU145-sh11.02 cells. Treatment with 3 Gy IR increased the apoptotic fraction of LNCaP cells from 3.53% to 6.90% (*p*=0.0174) and non-targeting siRNA treated LNCaP cells from 4.16% to 7.93% (*p*=0.0305). However, the most significant effect was observed in cells transfected with CXCR1/2-siRNA, where concurrent exposure to IR increased the apoptotic cell fraction from 4.20% to 21.76% (*p*=0.0097; Figure 4C). Similar results were obtained in DU145 sh11.02 cells, where the addition of IR increased the apoptotic fraction of CXCR1/2-siRNA treated cells from 17.66% to 27.70% (*p*=0.0071; Figure 4D).

Additional experiments were performed to assess the DNA-damage response following IR exposure and CXCR1/2-inhibition. Interestingly, PTEN-deficient sh11.02 cells had higher basal levels of 53BP1 foci and a slower rate of repair following exposure to 1 Gy IR compared to PTEN-expressing NT01 cells. However, attenuation of CXCR1/2-signaling had no additional effect on the DNA-damage response (Supplementary Figure 2).

### PTEN-loss and combined CXCR1/2-inhibition with radiation slows tumor growth in androgen-independent xenografts

Further experiments were conducted to evaluate the impact of CXCR1/2 inhibition upon the response of growing prostate tumours to radiotherapy. In these experiments, rather than use siRNA to deplete CXCR1/2 signaling potential, we employed CXCR1/2-targeted peptidomimetics termed “pepducins” which have been shown to uncouple the receptors from activating the intrinsic G proteins, blocking signal transduction in both *in vitro* and *in vivo* cancer models (Jamieson, Clarke et al., 2012). We validated the efficacy of the CXCR1/2-targeted pepducin, x1/2pal-i3, to attenuate CXCL-mediated tumorigenicity in *PTEN*-modulated prostate cancer cells, using an appropriate non-targeting peptide as the relevant control.

We initially validated the “pepducin” approach *in vitro*; as expected, using our DU145 NT01 and sh11.02 models, we observed that the addition of x1/2pal-i3 only increased radiation sensitivity *in vitro* in the PTEN-deficient context (Figure 5A; Supplementary Figure 3C). Experiments were extended *in vivo* following implantation of sh11.02 cells in the flank of SCID mice. Tumours (100 mm^3^) were subjected to daily injections of either the x1/2pal-i3 targeting peptide or the control peptide [x1/2pal-con] for five consecutive days and a single 2 Gy radiation exposure on day 2. On day 21, mice treated with the control pepducin (x1/2pal-con) displayed a mean tumor volume of 369.6±22.3 mm^3^. Treatment with x1/2pal-i3 alone resulted in a tumor growth delay after 21 days (226.1±16.0 mm^3^) which was comparable to mice treated with 2 Gy IR (249.9±15.9 mm^3^). However, the combined treatment consisting of 2 Gy IR and x1/2pal-i3 was most efficacious, resulting in a mean tumor volume of 166.6 ± 10.9 mm^3^ (Figure 5B). This combined therapy extended the median quadrupling time by 52% compared to tumours treated with IR tumors alone (*p*<0.001; Figure 5C). Treatment of PTEN-expressing NT01 tumours with x1/2pal-i3 offered no additive effect in combination with ionizing radiation (Supplementary Figure 3D).

**Figure 5.**
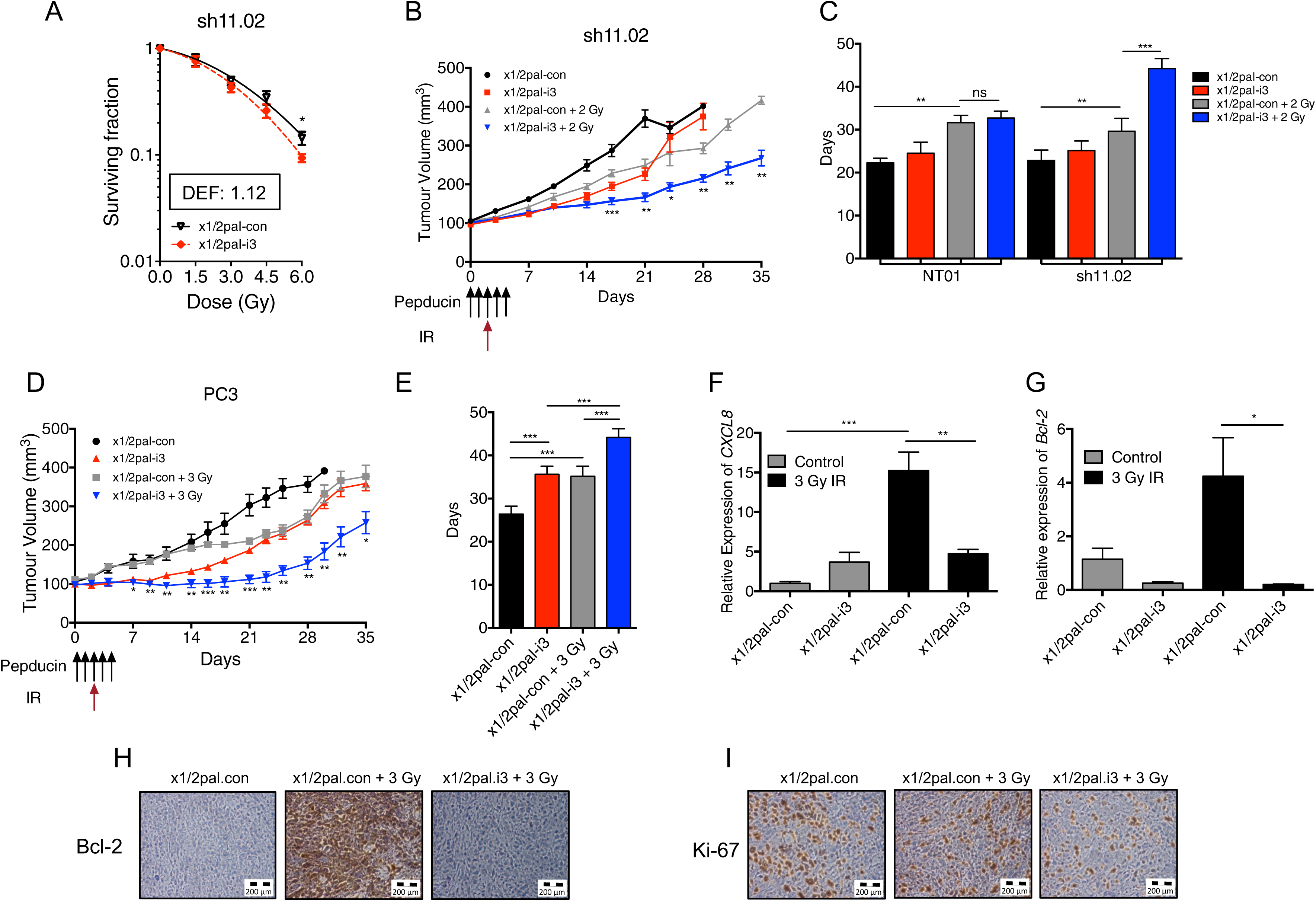
Inhibition of CXCR1/CXCR2-signaling delays tumor growth and potentiates radiotherapy-mediated growth delay of PTEN-null prostate cancer models. **A** Clonogenic survival curves showing DU145-sh11.02 survival fractions following treatment with increasing doses of IR. Cells were pre-treated 4H prior to IR exposure with either control pepducin (x1/2pal-con) or CXCR1/2-targeted pepducin (x1/2pal-i3). **B** Tumor growth curves showing sh11.02 xenograft tumor volumes. Mice were randomized into four groups (N=7/group): x1/2pal-con; x1/2pal-i3; x1/2pal-con + 2 Gy; and x1/2pal-ie + 2 Gy. Days in which pepducin (2 mg/kg) and IR treatments were performed are indicated. **C** Bar chart showing the mean time for PTEN-expressing NT01 and PTEN-deficient sh11.02 tumors to quadruple in size following treatment with the interventions modelled in (B). **D** Tumor growth curves showing PC3 xenograft tumor volumes. Mice were randomized into four groups (N=7/group): x1/2pal-con; x1/2pal-i3; x1/2pal-con + 3 Gy; and x1/2pal-ie + 3 Gy. Days in which pepducin (2 mg/kg) and IR treatments were performed are indicated. **E** Bar chart showing the mean time for PC3 tumors to quadruple in size following treatment with the interventions modelled in (D). **F-G** Bar charts showing gene expression of *CXCL8* and *BCL2* following treatment of PC3 xenografts with the interventions modelled in (D). Tumors were harvested on study day 5 before extracting RNA for qRT-PCR analysis. Both graphs show mice treated with pepducin but no IR (grey bars) or a combination of pepducin and IR (black bars). **H-I** Images depicting Bcl-2 and Ki-67 IHC analysis of PC3 xenografts harvested on study day 5. Scale bars indicate 200 μm. Data information: All data presented is in the format of mean ± SE. For (A) statistically significant differences for individual dose points were determined using t-tests. For tumor growth analysis statistically significant differences between radiation alone and combination treatment on specific study days were determined using t-tests (*p*<0.05*; *p*<0.01**; *p*<0.001***).

Further experiments were conducted in a second tumor model of *PTEN*-null PC3. As anticipated, administration of the x1/2pal-i3 pepducin increased sensitivity to IR and repressed CXCL8-induced Bcl-2 expression *in vitro*, while use of a non-targeting pepducin (x1/2pal-con) had no effect (Supplementary Figure 3A-B). Once again, we observed a profound effect of the CXCR1/2-targeting pepducin upon the growth of *PTEN*-null PC3 xenograft tumors. Relative to tumors treated with the control pepducin (x1/2pal-con; mean tumor volume of 303.0 ± 27.8 mm^3^ after 21 days), treatment with either IR (3 Gy) or the receptor-targeted x1/2pal-i3 pepducin produced a similar tumour growth delay with 21 day mean tumor volumes of 198.7±21.7 mm^3^ and 186.8±6.7 mm^3^, respectively. However, a combined use of radiation and administration of the x1/2pal-i3 pepducin attenuated tumor growth to a mean volume of 111.8±11.1 mm^3^ after 21 days (Figure 5D). Experiments were extended post-21 days and the tumor quadrupling time for each cohort was calculated (Figure 5E). There was no difference in tumor growth between the untreated control group (26.5±1.9 d) or those treated with the non-targeting x1/2pal-con peptide (26±1.3 d). Relative to x1/2pal-con-treated tumors, treatment with the CXCR1/2-targeting x1/2pal-i3 pepducin attenuated tumor growth (34.7±0.91 d; *p*=0.0003), producing a comparable growth delay as radiation (3 Gy; 35.7±2.008 d). Combined treatment of 3 Gy plus x1/2pal-i3 produced a mean tumor growth delay of 43±1.23 days, extending the time to reach the experimental endpoint by 20.4% over radiation alone. No acute toxicity was observed between the treatment groups as determined by changes in body weight in either the DU145 or PC3 models (Supplementary Figure 4A-C).

Pharmacodynamic markers were assessed in PC3 xenografted tumors harvested from additional mice within each experimental cohort at day five (48H post-IR). As predicted by our prior *in vitro* data, qRT-PCR analysis of the tumors confirmed that exposure to IR exposure elevated gene expression of *CXCL8* and *BCL2*; this induction of gene expression was attenuated in the presence of the receptor-targeted x1/2pal-i3 pepducin (*p*=0.005 and *p*=0.031 respectively; Figure 5F-G). Furthermore, IHC performed on tumor sections confirmed that IR exposure increased Bcl-2 in these *PTEN*-null tumors, which was again reversed by x1/2pal-i3-mediated inhibition of CXCR1/2-signaling (Figure 5H). Moreover, inhibition of CXCR1/2-signaling impaired the proliferative capacity of irradiated PC3 tumors as shown by reduction in the Ki-67 positive cell population (Figure 5I).

### The CXCR2-antagonist AZD5069 combined with radiation slows tumor growth in a PTEN-null xenograft model

To further validate our observations using a molecular-approach to abrogate CXCR2 signaling, we sought to use a pharmacological approach. AZD5069 is a selective, small molecule CXCR2 receptor antagonist that has shown tolerability in respiratory medicine conditions and is currently undergoing clinical evaluation in solid tumors, including advanced metastatic castrate-resistant prostate cancer (NCT03177187; “ACE” Trial) (De Soyza, Pavord et al., 2015). Accordingly, we used AZD5069 to determine whether a CXCR2-selective antagonist would phenocopy the impacts observed following administration of the receptor-targeting pepducin. C4-2 tumors were established in SCID mice and treatment with either vehicle or AZD5069 started when tumors reached a volume of 100 mm^3^. Radiation (3 Gy) was delivered on day 3 of this treatment regime. Combined therapy resulted in a tumor growth delay compared to either AZD5069 alone or 3 Gy IR alone (Figure 6A). Tumor quadrupling times were calculated to determine the benefit of combined therapy. Mice treated with vehicle control had a tumor quadrupling time of 6.33±2.04 days compared to 8.07±2.29 days in mice treated with AZD5069 (ns; *p*=0.181). Treatment with a single 3 Gy of IR resulted in a quadrupling time of 9.083±2.94 days. However, the greatest response was seen in mice exposed to combined therapy (16.92±3.31 days; *p*=0.0014), extending the time to reach experimental endpoint by 86.31% (Figure 6B). Importantly, no acute toxicity was observed between the treatment groups as determined by changes in body weight over the course of the experiment (Supplementary Figure 4D).

**Figure 6.**
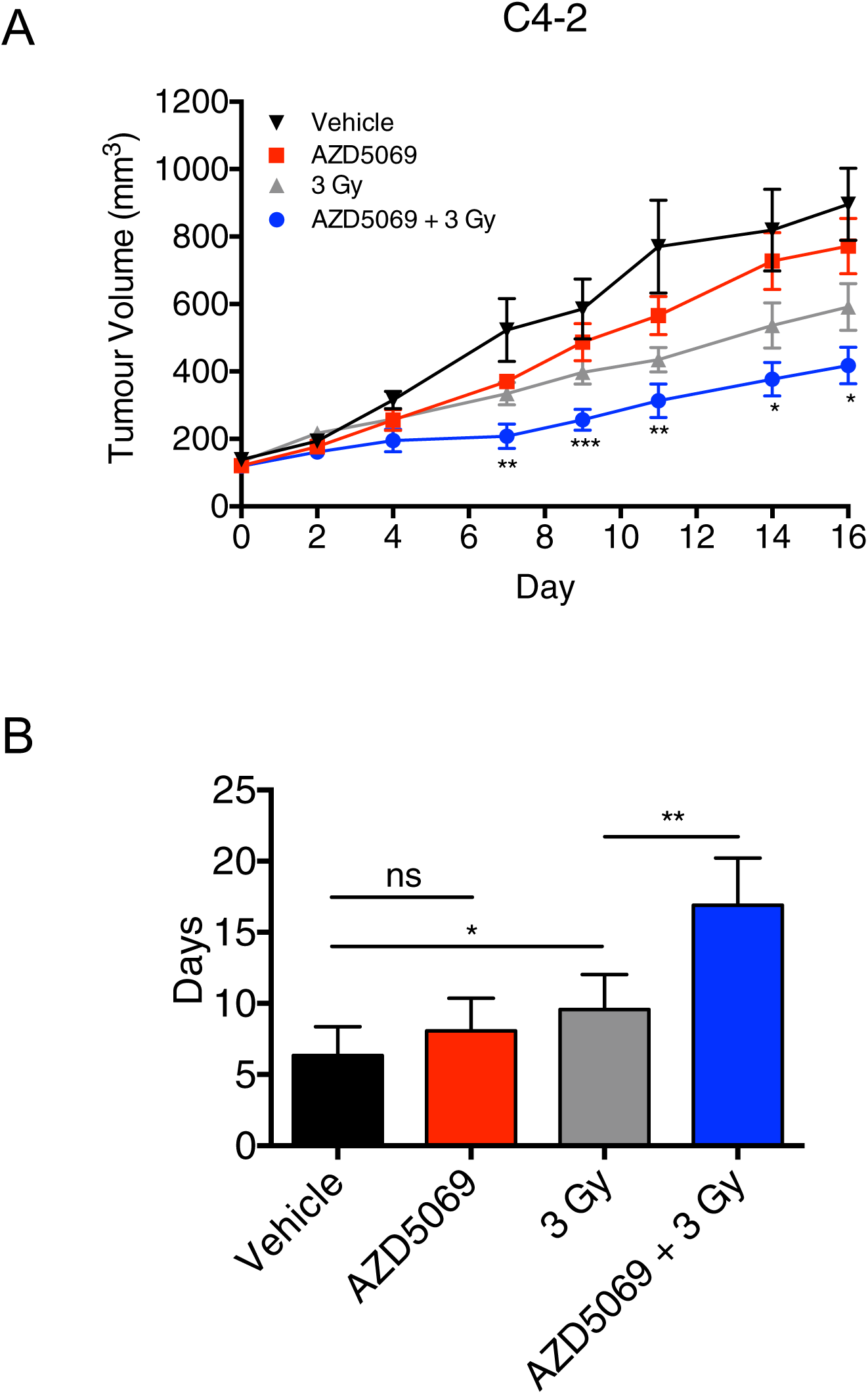
AZD5069-mediated CXCR2-inhibition increases C4-2 tumor radiosensitivity. **A** Tumor growth curves showing C4-2 xenograft tumor volumes. Mice were randomized into four groups (N=8/group): vehicle control; AZD5069; 3 Gy IR; and AZD5069 + 3 Gy IR. **B** Bar chart showing the mean time for C4-2 tumors to quadruple in size following treatment with the interventions modelled in (A). Data information: All data presented is in the format of mean±SE. For tumor growth analysis statistically significant differences between radiation alone and combination treatment on specific study days were determined using t-tests (*p*<0.05*; *p*<0.01**; *p*<0.001***).

## DISCUSSION

The use of radiation [external-beam radiotherapy in locally-advanced prostate cancer, stereotactic radiotherapy in the treatment of oligometastatic prostate cancer or radionuclides for resolution of bone disseminated CRPC] is a major treatment modality across the clinical landscape of prostate cancer. Technological advancements in the delivery and concentration of radioactivity to malignant zones of interest and use of devices to reduce radiation exposure to neighbouring tissues have greatly improved the response and tolerability of radiotherapy. However, over one-third of patients with organ-confined disease experience distant relapse while radionuclides can extend survival but not overcome the incurable aspect of metastatic disease (Agarwal, Sadetsky et al., 2008, Heidenreich, Gillessen et al., 2019). To further improve the efficacy of radiotherapy it is important to understand the biological mediators of radiotherapy resistance as foundational knowledge to characterise potential drug-radiotherapy combination regimens.

*PTEN* status has prognostic value in identifying patients at high risk of relapse post-radiotherapy (Zafarana et al., 2012a). However, biological drivers underpinning PTEN-associated relapse are yet to be characterised or evaluated clinically. We have previously shown that *PTEN*-loss selectively potentiates CXCL8 signaling in pre-clinical human and genetically-engineered murine models of prostate cancer (Maxwell et al., 2013). The association of PTEN status with elevated chemokine signaling was confirmed by analysis of both the MSKCC and the FASTMAN transcriptomic databases, which identified the presence of a PTEN^LOW^/ CXCR1/2^HIGH^ cluster, which correlated with increased rates of biochemical recurrence following either radical prostatectomy (MSKCC) or radical radiotherapy with curative intent (FASTMAN). Moreover, we were able to establish the relationship of this PTEN^LOW^/ CXCR1/2^HIGH^ cluster with distant metastasis in the FASTMAN cohort.

Our further experiments sought to establish a functional role of elevated chemokine signaling in the resistance of PTEN-deficient prostate cancer cells to radiotherapy. Our data confirms that exposure of PTEN-deficient but not PTEN-expressing cells to clinically-relevant doses of IR increases the expression of CXCL8, a chemokine that induces the activation of CXCR1 and CXCR2 receptors. Moreover, expression of both receptors was shown to increase in prostate cancer cell lines, albeit, independent of their PTEN status. Subsequent cell colony assays confirmed that the promotion of increased chemokine signaling was coupled to an adverse response of irradiated PTEN-deficient prostate cancer cells, and consequently, that the inhibition of this chemokine signaling pathway markedly increased the sensitivity of three distinct models of prostate carcinoma to IR. The use of isogenic cells with siRNA-, shRNA- or tetracycline-induced differential expression of PTEN confirmed that the abrogation of chemokine signaling only conferred radiosensitization in the context of *PTEN*-deficiency. Furthermore, a more pronounced effect of inhibiting chemokine signaling was observed at higher doses of IR, consistent with the current higher doses used in hypo-fractionated or stereotactic-radiotherapy protocols (Dearnaley et al., 2016).

Inhibition of CXCR1/2-signaling *in vitro* did not modulate the magnitude of DNA-foci formation or the rate of DNA repair in these cells. Instead, our data suggests that radiation-induced CXCR1/2-signaling sustains the viability of *PTEN*-depleted cells in the aftermath of radiation exposure. Specifically, inhibition of CXCR1/2-signaling prior to IR-exposure in *PTEN*-null LNCaP and *PTEN*-depleted DU145 cells significantly induced apoptosis, characterized by Annexin V/PI flow cytometry analysis and the induction of caspase- and PARP-cleavage. This is consistent with our prior studies confirming that inhibition of CXCR2-signaling potentiated oxaliplatin-induced apoptosis in PC3 cells (Wilson et al., 2008).

Our *in vivo* experimentation in three distinct prostate tumour models provides further compelling evidence for the contribution of CXCR1/CXCR2-signaling in adversely affecting radiotherapy outcome. Administration of a dual CXCR1/2-targeted antagonistic peptide alone inhibited the growth of *PTEN*-deficient but not *PTEN*-expressing DU145 and PC3 xenograft tumors, consistent with our prior and current demonstration of CXCL8 functioning as a key survival factor in this genetic context (Maxwell et al., 2013). Interestingly, the CXCR1/2-targeted pepducin was equi-effective in enabling an anti-tumor response as a clinically-relevant dose of radiation in both PTEN-depleted DU145 and PC3 tumours. However, of even greater significance, we observed that peptide-mediated inhibition extended the growth delay afforded by exposure of both *PTEN*-depleted DU145 and PC3 models to IR. These observations were further validated by experimentation in a third model, wherein the sensitivity of the LNCaP C4-2 to IR was increased by the administration of the small molecule antagonist of the CXCR2 receptor, AZD5069. Therefore, we have shown consistent responses across three distinct PTEN-deficient prostate cancer models, using both a molecular and pharmacological interventions.

The pronounced effect of targeting CXCR1 and CXCR2 signaling observed in our *in vivo* models is consistent with the multi-factorial role of chemokines within the tumour microenvironment. The baseline and radiation-enhanced secretion of CXCL8 (and its orthologues CXCL1, CXCL2, and CXCL5) from PTEN-deficient prostate tumor epithelial cells exerts both autocrine and paracrine actions within the tumor microenvironment given the expression of CXCR1 and CXCR2 upon multiple cell types (Acosta, O’Loghlen et al., 2008, Singh, Wu et al., 2011). Firstly, our *in vitro* data confirms a radiation-induced potentiation of CXCR1 and CXCR2 signaling in PTEN-deficient cells, which increases anti-apoptotic gene and protein expression. Analysis of pharmacodynamic markers within harvested PTEN-deficient tumour samples similarly indicated IR to induce the expression of Bcl-2, a known mediator of radiotherapy-resistance in a number of cancer models, but that this was attenuated by inhibition of CXCR1/CXCR2 signaling (An, Chervin et al., 2007). Secondly, tumor-derived CXCL signaling is likely to exert paracrine activation on surrounding stromal fibroblasts and monocyte-derived immune cells (Armstrong et al., 2016). Recent studies have confirmed that inhibition of CXCR2 signaling can repress the activity of myeloid-derived suppressor cells (MDSCs) and re-educate the differentiation of tumor-associated macrophages in genetically-engineered mouse models (Di Mitri, Mirenda et al., 2019). The broader significance of inhibiting CXCR-signaling upon the constitution and activity of the tumor microenvironment following exposure to radiotherapy is worthy of future targeted and more comprehensive investigation.

In conclusion, we have characterised a distinct cluster of primary prostate cancer defined by *PTEN*^LOW^, CXCR1^HIGH^ and CXCR2^HIGH^ expression that associates with adverse downstream clinical outcome. We have provided further experimental evidence using established models of disease that this biology may be a functional driver of the impaired radiotherapy response observed in PTEN-deficient tumours. Combined, our *in vitro* and *in vivo* data demonstrates that subjecting *PTEN*^LOW^ cancer cells or tumors to clinically-relevant doses of IR amplifies chemokine signaling, potentiating autocrine and paracrine signaling throughout the microenvironment that supports the survival of PTEN-deficient cells and facilitates the acquisition of a myeloid-enriched and potentially immunosuppressive microenvironment. Molecular-mediated and pharmacological-based inhibition of CXCR1/CXCR2-signaling sensitised each of the three PTEN-deficient tumour models to clinically relevant doses of IR. Consequently, we propose that targeting this chemokine signaling pathway may constitute a relevant therapeutic strategy to enhance the response of *PTEN*-deficient prostate carcinomas to radiation therapy. Given that functional impairment of *PTEN* is reported in up to 30% of all primary prostate cancer and that radical radiotherapy is a major treatment option for locally-advanced high-risk disease, CXCR1/CXCR2-targeted therapeutics may have a significant impact across a large cohort of patients.

Furthermore, the observation that chemokine receptor inhibition may have greater impacts at even higher radiation doses suggests that this intervention may have even greater significance in respect of the use of stereotactic radiotherapy in resolution of oligometastatic disease, and potentially, where there is further enrichment in the prevalence of *PTEN* gene aberration.

## Acknowledgements

This work was supported by grants from Cancer Research UK (A4106/C11512), the Medical Research Council (MR/J007641/1), the Movember/Prostate Cancer UK Centre of Excellence Award (CEO_13-002) and a Prostate Cancer UK Travelling Prize Fellowship (TLD-PF16-002).

The samples used in this research were received from the Northern Ireland Biobank which is funded by HSC Research and Development Division of the Public Health Agency in Northern Ireland and Cancer Research UK through the Belfast CR-UK Centre and the Northern Ireland Experimental Cancer Medicine Centre (ECMC); additional support was received from the Friends of the Cancer Centre. The Northern Ireland Molecular Pathology Laboratory which is responsible for creating resources for the NIB has received funding from Cancer Research UK, the Friends of the Cancer Centre and the Sean Crummey Foundation.

---

**Supplementary Table 1.** Western blot antibody information.

| Antibody | Supplier | Cat no. | Dilution |
| --- | --- | --- | --- |
| **Akt** | Cell Signaling | 9272 | 1:1000 |
| **pAkt (Ser473)** | Cell Signaling | 9271 | 1:1000 |
| **β-Actin** | Cell Signaling | 3700 | 1:5000 |
| **Bcl-2** | Santa Cruz | sc-7382 | 1:2500 |
| **Cleaved Caspase 9** | Cell Signaling | 7237 | 1:1000 |
| **CXCL8** | R&D Systems | MAB208 | 1:750 |
| **CXCR1** | R&D Systems | MAB330 | 1:750 |
| **CXCR2** | R&D Systems | MAB331 | 1:750 |
| **JAK2** | Abcam | ab108596 | 1:1000 |
| **pJAK2 (Tyr1007/8)** | Abcam | ab32101 | 1:1000 |
| **PARP** | Cell Signaling | 9542 | 1:1000 |
| **PTEN** | Cell Signaling | 9559 | 1:1000 |
| **STAT3** | Abcam | ab68153 | 1:1000 |
| **pSTAT3 (Tyr705)** | Abcam | ab76315 | 1:1000 |

**Supplementary Table 2.** Cell doubling times (± SEM) for *PTEN*-functional and ablated DU145 cells.

| Cell line | 0 Gy NT | 0 Gy<br>siCXCR1/2 | 3 Gy + siNT | 3 Gy<br>siCXCR1/2 |
| --- | --- | --- | --- | --- |
| **DU145 NT01** | 2.163 ± 0.022 | 2.118 ± 0.021<br>(□ ***) | 2.453 ± 0.025 | 4.627 ± 0.021<br>(□ ***) (⊠ ***) |
| **DU145 sh11.02** | 2.059 ± 0.041 | 2.148 ± 0.022<br>(α ***) | 2.238 ± 0.024<br>(⊠ ***) | 6.853 ± 0.033<br>(α ***) (⊠ ***)<br>(⊠ ***) |

Cell growth curves were established and monitored over a 10-day period. Populations were exposed to either sham-radiation (0 Gy) or radiation (3 Gy), in the absence (siNT) or presence (siCXCR1/2) of a CXCR1/CXCR2-targeted siRNA strategy.
Raw data was fit to an exponential growth equation (Prism 6 software) from which the mean doubling time for each cell population was calculated. Statistically-significant differences in cell doubling were determined using a two-way ANOVA.
□ represents a statistically-significant difference afforded by the provision of radiation to CXCR1/2-ablated, *PTEN*-expressing NT01-transfected cells.
α represents a statistically-significant difference afforded by the provision of radiation to CXCR1/2-ablated, *PTEN*-ablated sh11.02 cells.
⊠ represents a statistically-significant difference afforded by CXCR1/2-ablation in irradiated (3 Gy) *PTEN*-ablated sh11.02 cells ( $p < 0.0001$ ).
⊠ represents a statistically-significant difference afforded by combined radiation and CXCR1/2-ablation in *PTEN*-deficient cells ( $p < 0.0001$ ).

## Supplementary Figure Legends

**Supplementary Figure 1. Restoration of PTEN expression abrogates CXCR1/2-mediated radiosensitivity.**

**A-B** Clonogenic survival curves generated from PC3 and PC3+PTEN cells respectively. Cells were pre-treated with 25nM non-targeting siRNA (siNT) or CXCR1 and CXCR2 siRNA (siCXCR1/2) for 48H prior to IR treatment.

Data information: All data presented is a representation of N=3 independent experiments. All survival curves are fitted to a linear quadratic model and Dose Enhancement Factor (DEF) was calculated using the mean inactivation dose based on the area under the curve at a surviving fraction of 0.1. Statistically significant differences were determined using t-tests (*p*<0.05*; *p*<0.01**; *p*<0.001***).

**Supplementary Figure 2. Loss of PTEN expression impairs DNA damage repair but CXCR1/2 inhibition has no effect on the DNA damage response.**

**A** Immunofluorescence images showing 53BP1 foci formation (red) in DU145 NT01 and sh11.02 cells at several timepoints following treatment with 1 Gy IR. Nucleus staining (blue) is also shown via the use of DAPI.

**B** Bar chart showing the average number of foci per cell under control conditions in both NT01 and sh11.02 cells.

**C** Graph presenting the kinetic profile of double-strand break repair in *PTEN*-functional NT01 and *PTEN*-depleted sh11.02 DU145 cells over a 24H time-period following exposure to ionizing radiation (1 Gy).

**D-E** Bar charts showing average number of 53BP1 foci in NT01 and sh11.02 cells respectively under control conditions or 4H post IR treatment. Cells were pre-treated with non-targeting siRNA (black bars) or CXCR1/2 siRNA (grey bars) to determine the effect of CXCR1/2-inhibition on the DNA damage response.

Data information: All data presented is a representation of N=3 independent experiments. Statistically significant differences were determined using t-tests (*p*<0.05*; *p*<0.01**; *p*<0.001***). No significant difference is indicated using *ns*.

**Supplementary Figure 3. CXCR1/2-targeted pepducins in PTEN-null and PTEN-expressing prostate cancer cell lines.**

**A** Clonogenic survival curves showing PC3 survival fractions following treatment with increasing doses of IR. Cells were pre-treated 4H prior to IR exposure with either control pepducin (x1/2pal-con) or CXCR1/2-targeted pepducin (x1/2pal-i3).

**B** Bar chart showing *BCL2* gene expression under control conditions (grey bars) or following addition of 3nM recombinant human CXCL8 (black bars). Cells were pre-treated 4H prior to IR exposure with either control pepducin (x1/2pal-con) or CXCR1/2-targeted pepducin (x1/2pal-i3).

**C** Clonogenic survival curves showing DU145 NT01 survival fractions following treatment with increasing doses of IR. Cells were pre-treated 4H prior to IR exposure with either control pepducin (x1/2pal-con) or CXCR1/2-targeted pepducin (x1/2pal-i3).

**D** Tumor growth curves showing NT01 xenograft tumor volumes. Mice were randomized into four groups (N=7/group): x1/2pal-con; x1/2pal-i3; x1/2pal-con + 2 Gy; and x1/2pal-ie + 2 Gy. Days in which pepducin (2 mg/kg) and IR treatments were performed are indicated.

Data information: All data presented is in the format of mean ± SE. For (A) statistically significant differences for individual dose points were determined using t-tests. For tumor growth analysis statistically significant differences between radiation alone and combination treatment on specific study days were determined using t-tests (*p*<0.05*; *p*<0.01**; *p*<0.001***).

**Supplementary Figure 4. Body weight assessment following treatment with CXCR1/2-targeted agents alongside radiotherapy.**

**A-C** Graphs showing percentage body weight change relative to study day 0 in mice implanted with DU145 NT01, sh11.02 or PC3 cells respectively. Each graph shows the effect of the following treatments on body weight: x1/2pal-con; x1/2pal-i3; x1/2pal-con + 2 Gy IR; or x1/2pal-i3 + 2 Gy IR.

**D** Graph showing percentage body weight change in mice implanted with C4-2 cells following treatment with vehicle control, AZD5069, 3 Gy IR or AZD5069 + 3 Gy IR (Combination).

Data information: No significant difference was observed in any model as determined by ANOVA.

## Notes

**Disclosure of Potential Conflicts of Interest:** The authors declare that they have no conflict of interest.

## REFERENCES

Acosta JC, O’Loghlen A, Banito A, Guijarro MV, Augert A, Raguz S, Fumagalli M, Da Costa M, Brown C, Popov N, Takatsu Y, Melamed J, d’Adda di Fagagna F, Bernard D, Hernando E, Gil J (2008) Chemokine signaling via the CXCR2 receptor reinforces senescence. Cell 133: 1006–18

Agarwal PK, Sadetsky N, Konety BR, Resnick MI, Carroll PR, Cancer of the Prostate Strategic Urological Research E (2008) Treatment failure after primary and salvage therapy for prostate cancer: likelihood, patterns of care, and outcomes. Cancer 112: 307–14

An J, Chervin AS, Nie A, Ducoff HS, Huang Z (2007) Overcoming the radioresistance of prostate cancer cells with a novel Bcl-2 inhibitor. Oncogene 26: 652–61

Arcangeli G, Saracino B, Arcangeli S, Gomellini S, Petrongari MG, Sanguineti G, Strigari L (2017) Moderate Hypofractionation in High-Risk, Organ-Confined Prostate Cancer: Final Results of a Phase III Randomized Trial. J Clin Oncol 35: 1891–1897

Armstrong CW, Maxwell PJ, Ong CW, Redmond KM, McCann C, Neisen J, Ward GA, Chessari G, Johnson C, Crawford NT, LaBonte MJ, Prise KM, Robson T, Salto-Tellez M, Longley DB, Waugh DJ (2016) PTEN deficiency promotes macrophage infiltration and hypersensitivity of prostate cancer to IAP antagonist/radiation combination therapy. Oncotarget 7: 7885–98

Barker HE, Paget JT, Khan AA, Harrington KJ (2015) The tumour microenvironment after radiotherapy: mechanisms of resistance and recurrence. Nat Rev Cancer 15: 409–25

Catton CN, Lukka H, Gu CS, Martin JM, Supiot S, Chung PWM, Bauman GS, Bahary JP, Ahmed S, Cheung P, Tai KH, Wu JS, Parliament MB, Tsakiridis T, Corbett TB, Tang C, Dayes IS, Warde P, Craig TK, Julian JA et al. (2017) Randomized Trial of a Hypofractionated Radiation Regimen for the Treatment of Localized Prostate Cancer. J Clin Oncol 35: 1884–1890

Dai H, Wang C, Yu Z, He D, Yu K, Liu Y, Wang S MiR-17 Regulates Prostate Cancer Cell Proliferation and Apoptosis Through Inhibiting JAK-STAT3 Signaling Pathway.

Darwish OM, Raj GV (2012) Management of biochemical recurrence after primary localized therapy for prostate cancer. Front Oncol 2: 48

De Soyza A, Pavord I, Elborn JS, Smith D, Wray H, Puu M, Larsson B, Stockley R (2015) A randomised, placebo-controlled study of the CXCR2 antagonist AZD5069 in bronchiectasis. Eur Respir J 46: 1021–32

Dearnaley D, Syndikus I, Mossop H, Khoo V, Birtle A, Bloomfield D, Graham J, Kirkbride P, Logue J, Malik Z, Money-Kyrle J, O’Sullivan JM, Panades M, Parker C, Patterson H, Scrase C, Staffurth J, Stockdale A, Tremlett J, Bidmead M et al. (2016) Conventional versus hypofractionated high-dose intensity-modulated radiotherapy for prostate cancer: 5-year outcomes of the randomised, non-inferiority, phase 3 CHHiP trial. Lancet Oncol 17: 1047–1060

Di Mitri D, Mirenda M, Vasilevska J, Calcinotto A, Delaleu N, Revandkar A, Gil V, Boysen G, Losa M, Mosole S, Pasquini E, D’Antuono R, Masetti M, Zagato E, Chiorino G, Ostano P, Rinaldi A, Gnetti L, Graupera M, Martins Figueiredo Fonseca AR et al. (2019) Re-education of Tumor-Associated Macrophages by CXCR2 Blockade Drives Senescence and Tumor Inhibition in Advanced Prostate Cancer. Cell Rep 28: 2156–2168 e5

Ferraldeschi R, Nava Rodrigues D, Riisnaes R, Miranda S, Figueiredo I, Rescigno P, Ravi P, Pezaro C, Omlin A, Lorente D, Zafeiriou Z, Mateo J, Altavilla A, Sideris S, Bianchini D, Grist E, Thway K, Perez Lopez R, Tunariu N, Parker C et al. (2015) PTEN protein loss and clinical outcome from castration-resistant prostate cancer treated with abiraterone acetate. Eur Urol 67: 795–802

Heidenreich A, Gillessen S, Heinrich D, Keizman D, O’Sullivan JM, Carles J, Wirth M, Miller K, Reeves J, Seger M, Nilsson S, Saad F (2019) Radium-223 in asymptomatic patients with castration-resistant prostate cancer and bone metastases treated in an international early access program. BMC Cancer 19: 12

Jain S, Lyons CA, Walker SM, McQuaid S, Hynes SO, Mitchell DM, Pang B, Logan GE, McCavigan AM, O’Rourke D, McArt DG, McDade SS, Mills IG, Prise KM, Knight LA, Steele CJ, Medlow PW, Berge V, Katz B, Loblaw DA et al. (2018) Validation of a Metastatic Assay using biopsies to improve risk stratification in patients with prostate cancer treated with radical radiation therapy.

Jamieson T, Clarke M, Steele CW, Samuel MS, Neumann J, Jung A, Huels D, Olson MF, Das S, Nibbs RJ, Sansom OJ (2012) Inhibition of CXCR2 profoundly suppresses inflammation-driven and spontaneous tumorigenesis. J Clin Invest 122: 3127–44

Khemlina G, Ikeda S, Kurzrock R (2015) Molecular landscape of prostate cancer: implications for current clinical trials. Cancer Treat Rev 41: 761–6

Kumareswaran R, Ludkovski O, Meng A, Sykes J, Pintilie M, Bristow RG (2012) Chronic hypoxia compromises repair of DNA double-strand breaks to drive genetic instability. J Cell Sci 125: 189–99

Maxwell PJ, Coulter J, Walker SM, McKechnie M, Neisen J, McCabe N, Kennedy RD, Salto-Tellez M, Albanese C, Waugh DJ (2013) Potentiation of inflammatory CXCL8 signalling sustains cell survival in PTEN-deficient prostate carcinoma. Eur Urol 64: 177–88

Maxwell PJ, Gallagher R, Seaton A, Wilson C, Scullin P, Pettigrew J, Stratford IJ, Williams KJ, Johnston PG, Waugh DJ (2007) HIF-1 and NF-kappaB-mediated upregulation of CXCR1 and CXCR2 expression promotes cell survival in hypoxic prostate cancer cells. Oncogene 26: 7333–45

McCabe N, Hanna C, Walker SM, Gonda D, Li J, Wikstrom K, Savage KI, Butterworth KT, Chen C, Harkin DP, Prise KM, Kennedy RD (2015) Mechanistic Rationale to Target PTEN-Deficient Tumor Cells with Inhibitors of the DNA Damage Response Kinase ATM. Cancer Res 75: 2159–65

Miyake M, Tanaka N, Asakawa I, Morizawa Y, Anai S, Torimoto K, Aoki K, Yoneda T, Hasegawa M, Konishi N, Fujimoto K (2014) Proposed salvage treatment strategy for biochemical failure after radical prostatectomy in patients with prostate cancer: a retrospective study. Radiat Oncol 9: 208

Muldermans JL, Romak LB, Kwon ED, Park SS, Olivier KR (2016) Stereotactic Body Radiation Therapy for Oligometastatic Prostate Cancer. Int J Radiat Oncol Biol Phys 95: 696–702

Murphy C, McGurk M, Pettigrew J, Santinelli A, Mazzucchelli R, Johnston PG, Montironi R, Waugh DJ (2005) Nonapical and cytoplasmic expression of interleukin-8, CXCR1, and CXCR2 correlates with cell proliferation and microvessel density in prostate cancer. Clin Cancer Res 11: 4117–27

Ost P, Jereczek-Fossa BA, As NV, Zilli T, Muacevic A, Olivier K, Henderson D, Casamassima F, Orecchia R, Surgo A, Brown L, Tree A, Miralbell R, De Meerleer G (2016) Progression-free Survival Following Stereotactic Body Radiotherapy for Oligometastatic Prostate Cancer Treatment-naive Recurrence: A Multi-institutional Analysis. Eur Urol 69: 9–12

Patel PH, Chaw CL, Tree AC, Sharabiani M, van As NJ (2019) Stereotactic body radiotherapy for bone oligometastatic disease in prostate cancer. World J Urol

Phin S, Moore MW, Cotter PD (2013) Genomic Rearrangements of PTEN in Prostate Cancer. Front Oncol 3: 240

Resnick MJ, Koyama T, Fan KH, Albertsen PC, Goodman M, Hamilton AS, Hoffman RM, Potosky AL, Stanford JL, Stroup AM, Van Horn RL, Penson DF (2013) Long-term functional outcomes after treatment for localized prostate cancer. N Engl J Med 368: 436–45

Singh S, Wu S, Varney M, Singh AP, Singh RK (2011) CXCR1 and CXCR2 silencing modulates CXCL8-dependent endothelial cell proliferation, migration and capillary-like structure formation. Microvasc Res 82: 318–25

Suzuki H, Freije D, Nusskern DR, Okami K, Cairns P, Sidransky D, Isaacs WB, Bova GS (1998) Interfocal heterogeneity of PTEN/MMAC1 gene alterations in multiple metastatic prostate cancer tissues. Cancer Res 58: 204–9

Taiakina D, Dal Pra A, Bristow RG (2014) Intratumoral hypoxia as the genesis of genetic instability and clinical prognosis in prostate cancer. Adv Exp Med Biol 772: 189–204

Taylor BS, Schultz N, Hieronymus H, Gopalan A, Xiao Y, Carver BS, Arora VK, Kaushik P, Cerami E, Reva B, Antipin Y, Mitsiades N, Landers T, Dolgalev I, Major JE, Wilson M, Socci ND, Lash AE, Heguy A, Eastham JA et al. (2010) Integrative genomic profiling of human prostate cancer. Cancer Cell 18: 11–22

Wilson C, Purcell C, Seaton A, Oladipo O, Maxwell PJ, O’Sullivan JM, Wilson RH, Johnston PG, Waugh DJ (2008) Chemotherapy-induced CXC-chemokine/CXC-chemokine receptor signaling in metastatic prostate cancer cells confers resistance to oxaliplatin through potentiation of nuclear factor-kappaB transcription and evasion of apoptosis. J Pharmacol Exp Ther 327: 746–59

Zafarana G, Ishkanian AS, Malloff CA, Locke JA, Sykes J, Thoms J, Lam WL, Squire JA, Yoshimoto M, Ramnarine VR, Meng A, Ahmed O, Jurisca I, Milosevic M, Pintilie M, van der Kwast T, Bristow RG (2012a) Copy number alterations of c-MYC and PTEN are prognostic factors for relapse after prostate cancer radiotherapy. Cancer 118: 4053–4062

