## Supplementary figures and images for "Targeting CXCR1 and CXCR2 to overcome radiotherapy resistance in PTEN-deficient prostate carcinoma"

### Supplementary Figures

Supplementary Figure 1

A

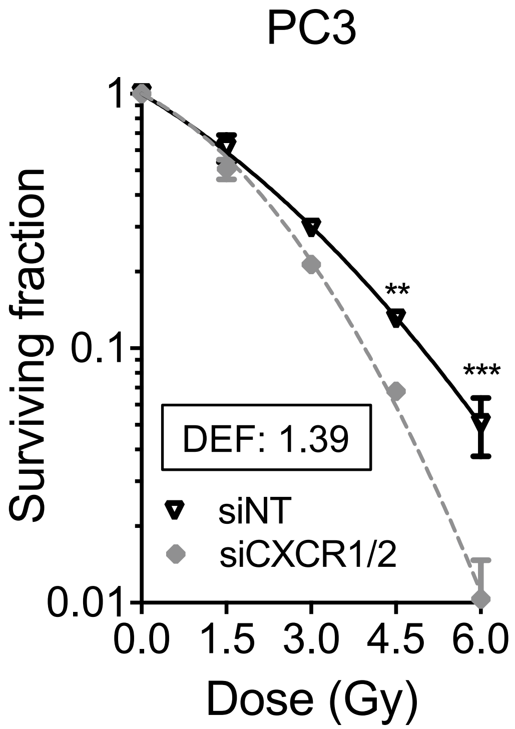


B

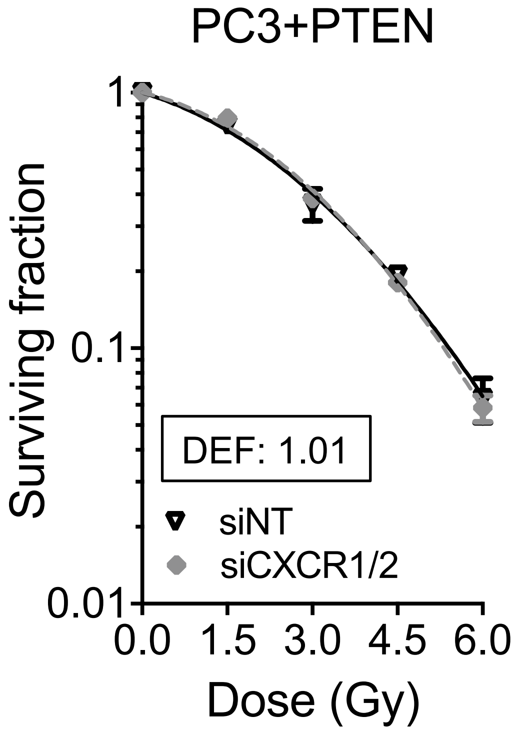

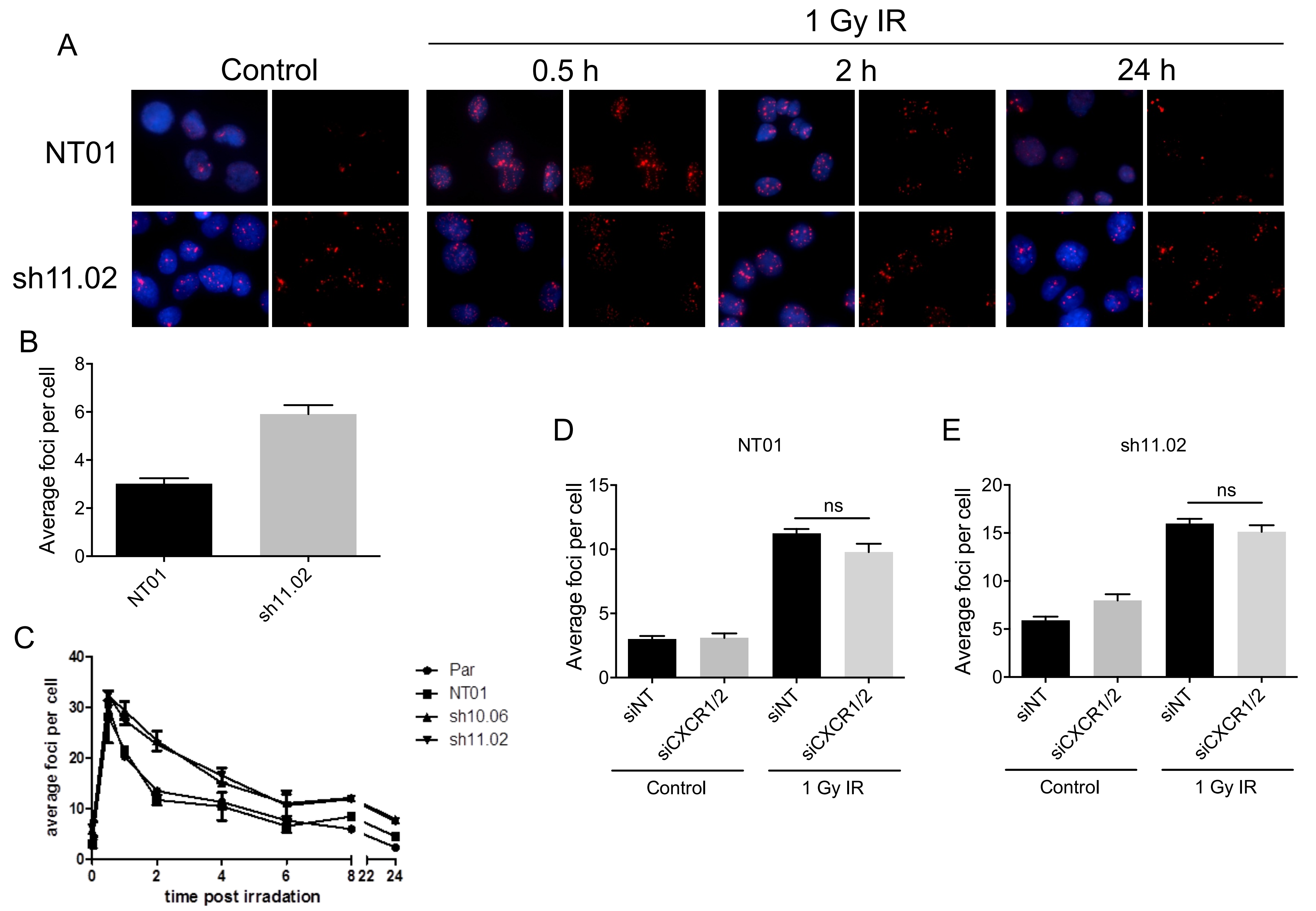

Supplementary Figure 3

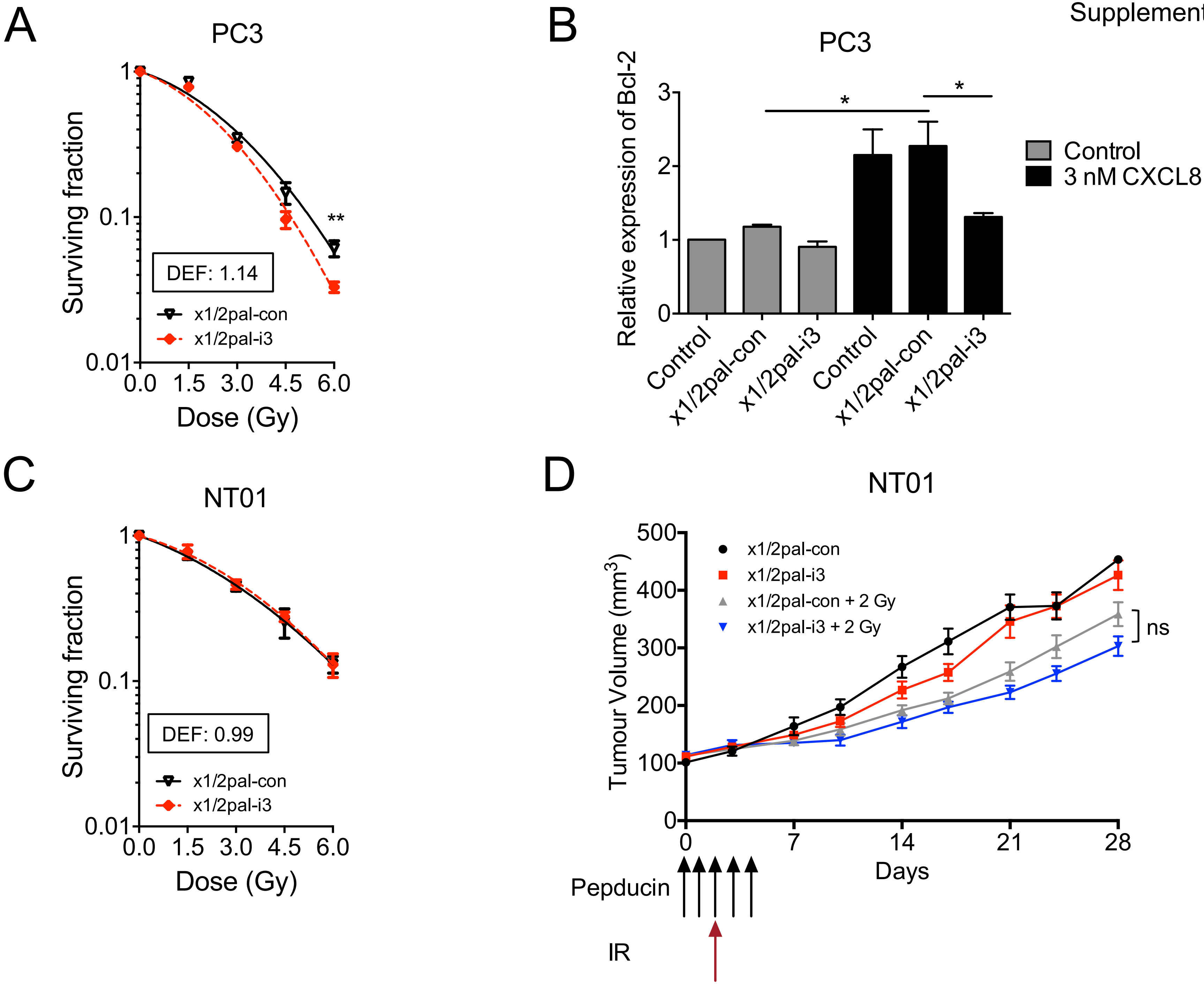

#### Supplementary Figure 4

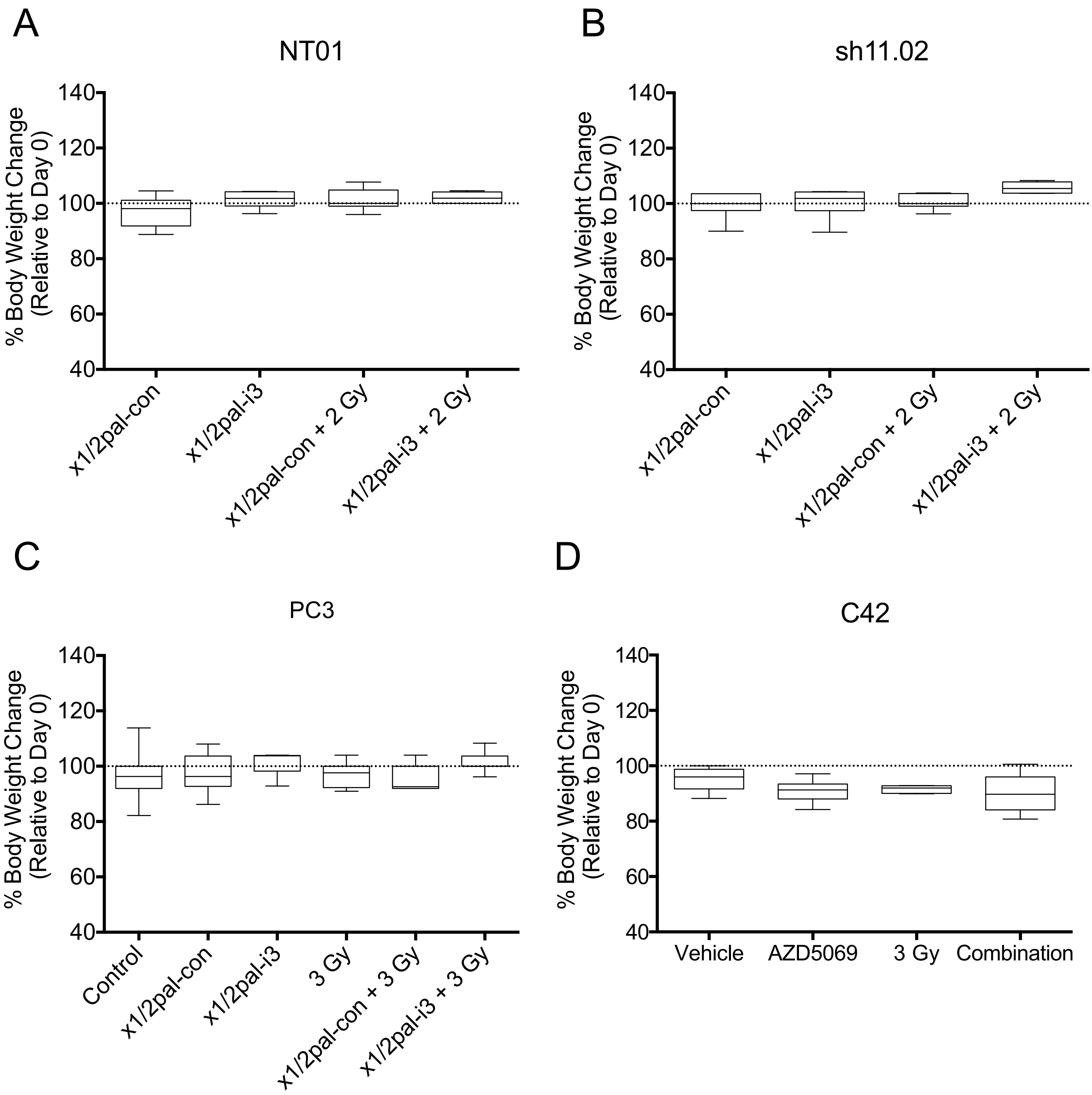
